# Auditory attention reorganizes the phase alignment of neural oscillations

**DOI:** 10.64898/2026.03.24.714038

**Authors:** Adi Korisky, Blair Kaneshiro, Radhika S Gosavi, Elizabeth Y Toomarian, Madison Bunderson, Bruce D McCandliss

## Abstract

Auditory attention enables the selection of behaviorally relevant sounds in dynamic environments, supporting the flexible allocation of neural resources over time. Although neural entrainment has been proposed as a mechanism for temporal prediction in audition, human studies have largely emphasized changes in response strength, leaving unresolved whether attention reorganizes the temporal alignment of entrained activity across hierarchical cortical networks. Here, we introduce the Selective Temporal Alignment of Components (STAC) framework to dissociate stimulus-driven and attention-controlled dynamics using non-invasive EEG. In a series of experiments across two independent adolescent cohorts (n = 79), Reliable Components Analysis (RCA) revealed two dissociable entrained networks with distinct spatial, functional, and attentional profiles: a sensory-driven network that remained tightly stimulus-locked and a frontal-auditory network that exhibited systematic attention-dependent phase shifts. These phase dynamics were consistent across independent cohorts and stable within individuals, and critically, predicted performance on a standardized neuropsychological measure of auditory attention. Together, these findings establish selective temporal alignment as a robust and behaviorally relevant neural mechanism underlying auditory attentional control.

## Introduction

More than 150 years ago, Hermann von Helmholtz proposed that attention plays a critical role in shaping the temporal dynamics of perception. In psychophysiological reaction-time experiments, he showed that temporal processing is stable within individuals yet sensitive to attentional state, noting that *“If at the time of perceiving the signal the thoughts are occupied with something else,…. it [the reaction] takes much more time”* (Helmholtz, 1850; as discussed in Schmidgen, 2002). This observation marked an early articulation of a core idea that continues to shape cognitive neuroscience - attention modulates *when* neural processes unfold, not only *what* information is processed.

Indeed, understanding how attention shapes the timing of neural processing remains a central goal in contemporary neuroscience, with extensive work examining how temporal and spectral features guide attentional selection in multisensory contexts. Among these, neural entrainment has emerged as a prominent framework for understanding the dynamic relationship between endogenous rhythmic brain activity, attention, and temporal prediction (Calderone et al., 2014; Gibbings et al., 2023; Gomez-Ramirez et al., 2011; Keitel et al., 2011; Nozaradan et al., 2011; Rimmele et al., 2018; Saupe et al., 2009; Schroeder & Lakatos, 2009). Broadly defined, neural entrainment refers to the synchronization of endogenous brain oscillations with the rhythmic structure of external stimuli, aligning both the phase (timing) and amplitude (strength) of neural activity with sensory input features (Lakatos et al., 2019; Obleser & Kayser, 2019; Schroeder & Lakatos, 2009). According to this view, when attention is directed toward a specific stream, top-down mechanisms are thought to bias this alignment, ensuring that the most acoustically relevant features of the target stimuli coincide with optimal phases of neural excitability, while irrelevant distractors are more likely to fall outside these windows and therefore receive reduced processing (Besle et al., 2011; Herbst & Landau, 2016; Lakatos et al., 2008, 2016; Obleser & Kayser, 2019; Schroeder & Lakatos, 2009; Thut et al., 2011; VanRullen, 2016; Vanrullen et al., 2011).

Within the auditory domain, how neural activity dynamically tracks the temporal structure of acoustic input is fundamental: unlike vision, which allows prioritizing information through spatial orienting, auditory perception relies heavily on timing to segregate acoustic streams and extract relevant information. Converging evidence demonstrates that in multisensory environments, attention modulates neural entrainment toward a target auditory stream, providing a mechanism that supports perceptual segmentation, enhances temporal prediction, and facilitates the efficient encoding of dynamic acoustic information across both non-verbal (Gomez-Ramirez et al., 2011; Kachlicka et al., 2022; Laffere et al., 2021; Nozaradan et al., 2011, 2012; Oever et al., 2017; Saupe et al., 2009) and verbal stimuli (Bree et al., 2021; Ding & Simon, 2014; Giraud & Poeppel, 2012; Har-shai Yahav & Zion Golumbic, 2021; Horton et al., 2013; Zion Golumbic et al., 2013; Zoefel & VanRullen, 2016). However, despite extensive research, most studies have focused on whether neural activity tracks attended rhythms more strongly, rather than on how attention reshapes the neural temporal dynamics of this tracking over time.

By contrast, approaches that have moved beyond strength-based metrics of auditory entrainment have largely relied on invasive intracranial electrophysiology, including work in non-human primates (Lakatos et al., 2008; Schroeder & Lakatos, 2009) and humans (Auksztulewicz et al., 2018; Besle et al., 2011). These studies have shown that the temporal operation of neural entrainment is divided into two distinct, phase-dependent levels of processing. The first level reflects fast sensory-driven entrainment arising from primary auditory regions, where neural activity closely tracks stimulus timing with a consistent cortical delay. The second level reflects attentional modulation, in which neural synchronization extends beyond primary sensory areas and the dominant oscillatory phase becomes governed by attention rather than external input solely. However, translating this two-level framework to broad human research using non-invasive methods has proven challenging. At the sensor level, it is difficult to disentangle overlapping neural contributions, as each frequency band typically exhibits a single dominant phase that may reflect a combination of sensory-driven and attention-modulated activity. This presents a challenge for studies using scalp electroencephalography (EEG), as the recorded phase at some sensors may be dominated by sensory-phase dynamics rather than attention-dependent modulation, potentially contributing to inconsistent results and interpretations across studies (Cabral-Calderin & Henry, 2022; Haegens & Zion Golumbic, 2018). As a result, the precise temporal processes through which auditory rhythms are transformed into neural oscillations in the temporal domain, and how attention modulates this transformation, remain poorly understood.

Motivated by this gap, in the current study, we pursue two primary aims. First, we examine how attention shapes the temporal dynamics of neural responses toward auditory input. We introduce the **Selective Temporal Alignment of Components (STAC)** approach, a methodological framework that dissociates stimulus-driven and attention-controlled dynamics, offering a non-invasive human analogue of two-level models described in animal work. The term *Selective* reflects the core principle of selective attention: prioritizing task-relevant input over competing stimuli. *Temporal Alignment* captures the central mechanism we propose—attention does not merely modulate the strength of neural responses but dynamically adjusts when neural activity aligns with rhythmic input, expressed as systematic phase shifts in entrained activity. Finally, *Components* refers to spatially dissociable processing levels identified through Reliable Components Analysis (RCA), reflecting separable neural sources consistent with hierarchical sensory-driven and attention-controlled stages.

To this end, we conducted two complementary EEG experiments to explore how selective attention modulates phase alignment to rhythmic auditory stimuli across sensory and attentional hierarchies, and how these dynamics manifest across late childhood and early adolescence. *Experiment 1* included children aged 10-12 years and examined how attention modulates the phase of the auditory steady-state response within different spatial components. Participants were simultaneously presented with a 3-Hz, speech-like auditory stream and a 1.25-Hz visual stream; they were required to selectively shift their attention between these two rhythmic inputs. We then applied RCA to spatially dissociate neural processing stages engaged during steady-state auditory stimulation. Prior work from our lab using the steady-state visual evoked potentials paradigm has shown that RCA can separate multiple components with distinct scalp topographies and dominant phase characteristics, each consistent across harmonics and differentially sensitive to cognitive manipulations (Wang et al., 2023, 2025). By extending this approach to the auditory domain, we first aimed to identify components with distinct cortical time courses, phase properties, and attentional sensitivity, and to test whether attentional modulation preferentially affects specific processing stages. *Experiment 2,* conducted with adolescents aged 12–14 years, served as a replication study, implementing the same design and analytic framework in an independent cohort to assess the reliability and behavioral relevance of the attentional modulation effects identified in *Experiment 1*.

## Results

### Behavioral Performance Confirms Effective Manipulation of Selective Attention

In both experiments, participants were presented with steady-state auditory (3 Hz syllable stream) and visual (1.25 Hz stream) stimuli. Depending on the condition, they were instructed to selectively attend to one modality and detect infrequent target events in the attended stream while ignoring the other modality.

In **Experiment 1,** a paired-samples t-test revealed significant differences between conditions across trials. Participants showed higher accuracy rates (t_(42)_ = 6.58, p <.0001, 95% CI [0.08, 0.16]) and lower reaction times (t_(42)_ = 4.61, p <.0001, 95% CI [-44.3,-17.3]) in the *Auditory-attended* condition, suggesting that the auditory target-detection was less challenging for participants compared to visual detection.

**Experiment 2** was composed of two sessions, and we conducted a 2×2 repeated-measures ANOVA (Condition × Session) on both accuracy rates and response times. Analysis of the accuracy rates revealed a significant main effect of Condition [*F*_(1,35)_ = 129.07, *p* <.001], showing that participants were significantly less accurate in the *Auditory-attended* compared to the *Auditory-ignored* condition. There was no significant main effect of Session [F_(1,35)_ = 2.66, p =.112] and no significant Condition × Session interaction [F_(1,35)_ = 1.66, p =.206], indicating that the performance difference between conditions remained stable across sessions and did not vary with practice or learning. For reaction times, results revealed a significant main effect of Condition [F_(1,35)_ = 72.52, p <.001], with responses significantly slower in the *Auditory-attended* compared with the *Auditory-ignored* condition. We also observed a significant interaction effect [F_(1,35)_ = 8.89, p =.005]. Post-hoc paired t-tests showed that although RTs were generally slower for auditory target detection, reaction times in the Auditory-attended condition decreased from Session 1 to Session 2, whereas in the Auditory-ignored condition, RTs increased between the sessions. Together, these results suggest that practice effects differed across conditions, with improvements emerging only when participants were required to focus on the auditory stream. No significant effect was observed for Session [F_(1,35)_ = 0.3, p =.58].

As described in the Methods, while Experiments 1 and 2 shared a similar auditory stream and identical frequency presentation rates, they differed in visual stimuli and several procedural aspects of the task. Consequently, despite the appeal of comparing the behavioral outcomes, especially given their opposing directions, the results are not directly comparable and should be interpreted independently within the context of each experiment.

### RCA Reveals Two Hierarchical Auditory Processing Components

As a first step of the STAC analysis, we applied RCA to dissociate hierarchical networks underlying auditory processing. RCA was conducted on the 3-Hz filtered data (and its harmonics) from the *Auditory-attended* and *Auditory-ignored* conditions. In both **Experiment 1** and **Experiment 2**, analysis revealed two reliable components, both of which showed significant eigenvalue coefficients as confirmed by phase-shuffling permutation testing (p < 0.001). In **Experiment 1,** the first reliable component (RC1) exhibited a broad central-frontal distribution centered around the midline (Across-trial covariates: dGen = 0.4) and a second component (RC2) displayed a more symmetrical anterior midline topography with bilateral temporal-parietal activation, slightly stronger for the left side (Across-trial covariates: dGen = 0.09). Similar to Experiment 1, in **Experiment 2,** the first component exhibited a broad central-frontal distribution but with a left-lateralized peak (Across trials covariance: dGen_(Session 1)_ = 0.42; dGen_(Session 2)_ = 0.36). The second component (RC2) displayed a more symmetrical anterior midline topography with two bilateral peaks over temporal-parietal sensors (Across trials covariance: dGen_(Session 1)_ = 0.19; dGen_(Session 2)_ = 0.22). See Figure 1 for all RC topographies.

**Figure 1:**
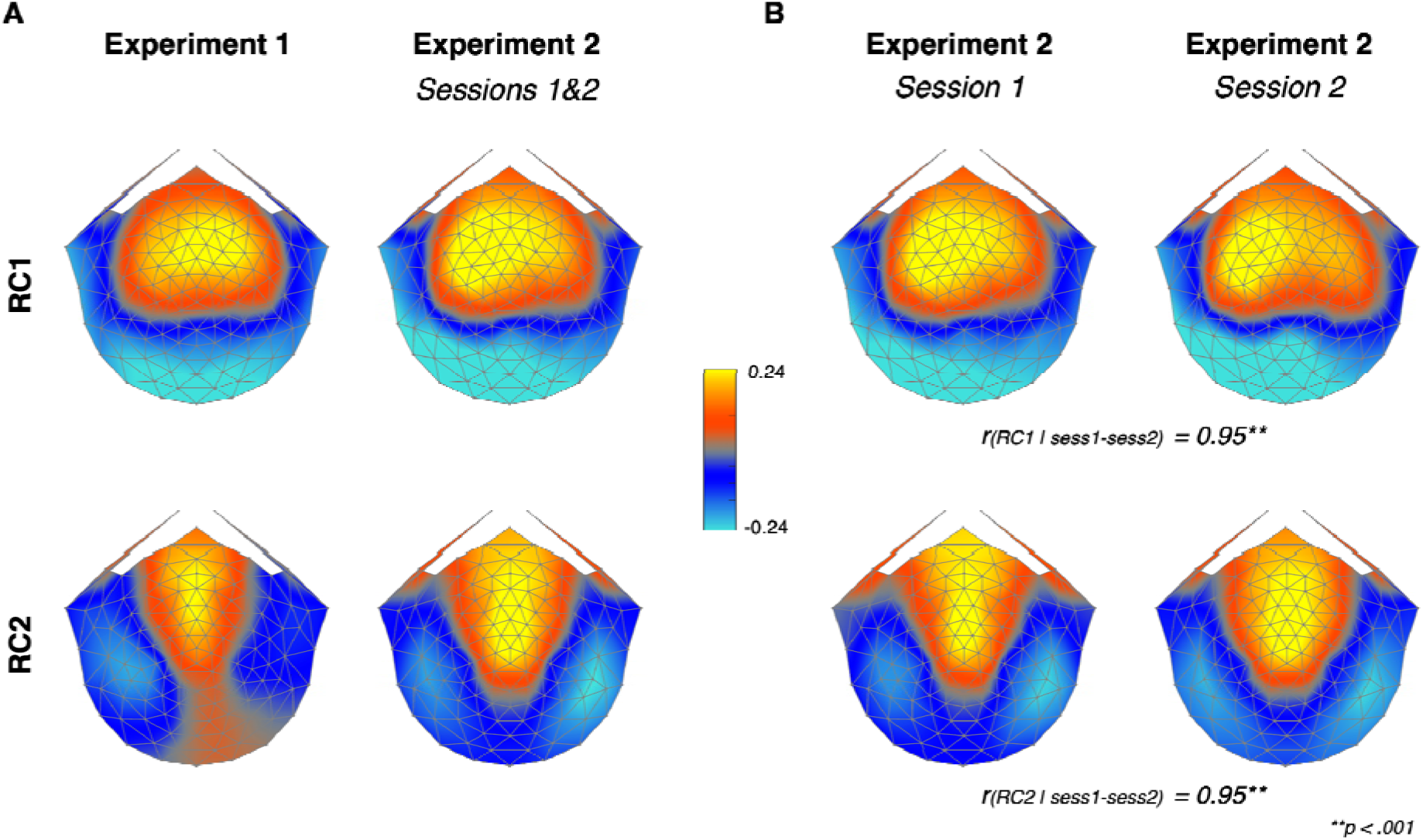
Topographic maps of the first two reliable components (RC1, top row; RC2, bottom row) across the two experiments. Forward-model–projected topographies (A-matrix) were computed from the filtered data (3-Hz and its harmonics). **(A)** Reliable components from Experiment 1 (left column) and Experiment 2 (right column, both sessions combined). **(B)** Reliable components from Experiment 2 are shown separately for the two sessions—Session 1 (left) and Session 2 (right). R values indicate the Pearson’s correlation of the computed A-matrix values across sessions.

Next, we conducted a Pearson correlation analysis comparing the forward-projected activity (columns of the *A* matrices) from the first and second sessions of Experiment 2. Results showed strong consistency across sessions, with robust positive correlations for RC1 (*r* = 0.95, *p* <.0001) and RC2 (*r* = 0.95, *p* <.0001). Given this high consistency, the two datasets were combined in all following statistical analyses to provide an integrated overview of the observed effects. All preprocessing and RCA computations were nonetheless conducted separately for each session as mentioned in the Methods section, to preserve the internal structure of each dataset.

### Attention Differentially Reorganizes Component-Level Temporal Dynamics

Next, to examine how auditory attention differentially modulates the temporal dynamics of the neural processes captured by each RC, we performed a paired-sample Hotelling test for circular data on the 3-Hz phase values derived from the component-space data, testing the main effects and the interaction (RC × Condition) between Component (RC1 vs. RC2) and Condition (*Auditory-attended vs. Auditory-ignored*). As mentioned in the Methods, because Experiments 1 and 2 differed in task and some experimental parameters, the results are not intended to be compared directly across experiments; instead, we focus on converging patterns and shared trends observed within each experiment.

In **Experiment 1,** a significant main effect was found for RC [F_(2, 41)_ = 10.95, *p* <.0005, Bonferroni corrected], showing that 3-Hz phases of RC1 data were centered around smaller phase values compared to those from RC2. Importantly, we found a significant interaction effect between Component and Condition [*F*_(2, 41)_ = 8.25, *p* <.005], showing that the influence of attention on the neural entrainment toward the 3-Hz auditory stream differed between components. Post-hoc paired-sample Hotelling’s tests revealed a significant attentional modulation effect in the RC2 component space data (F_(2, 41)_ = 3.57, p <.05), but not in RC1 [F_(2, 41)_ = 0.83, p =.40]. No other differences were found, and the main effect for condition was not significant [F_(2, 41)_ = 0.6, *p* = n.s].

Similar patterns were observed in **Experiment 2**. Our analysis revealed a significant main effect suggesting that 3-Hz phases derived from RC1 differ significantly from those derived by RC2 [*F*_(2, 70)_ = 24.8, *p* <.001]. Results also indicate a significant main effect for condition [F_(2, 70)_ = 13.7, *p* <.001], suggesting that when participants paid attention to the auditory stream, the 3-Hz phases in both components were smaller compared to those in the condition where participants actively ignored the audio. As in Experiment 1, we also found a significant interaction effect between component and condition [*F*_(2, 70)_ = 13.4, *p* <.001]. Post-hoc paired-sample Hotelling’s tests revealed a significant attentional modulation effect in the RC2 component space data [F_(2, 70)_ = 9.3, p <.001], but not in RC1 [F_(2, 70)_ = 2.2, p =.10]. See Figure 2 for full component-space data distributions and corresponding statistics.

**Figure 2:**
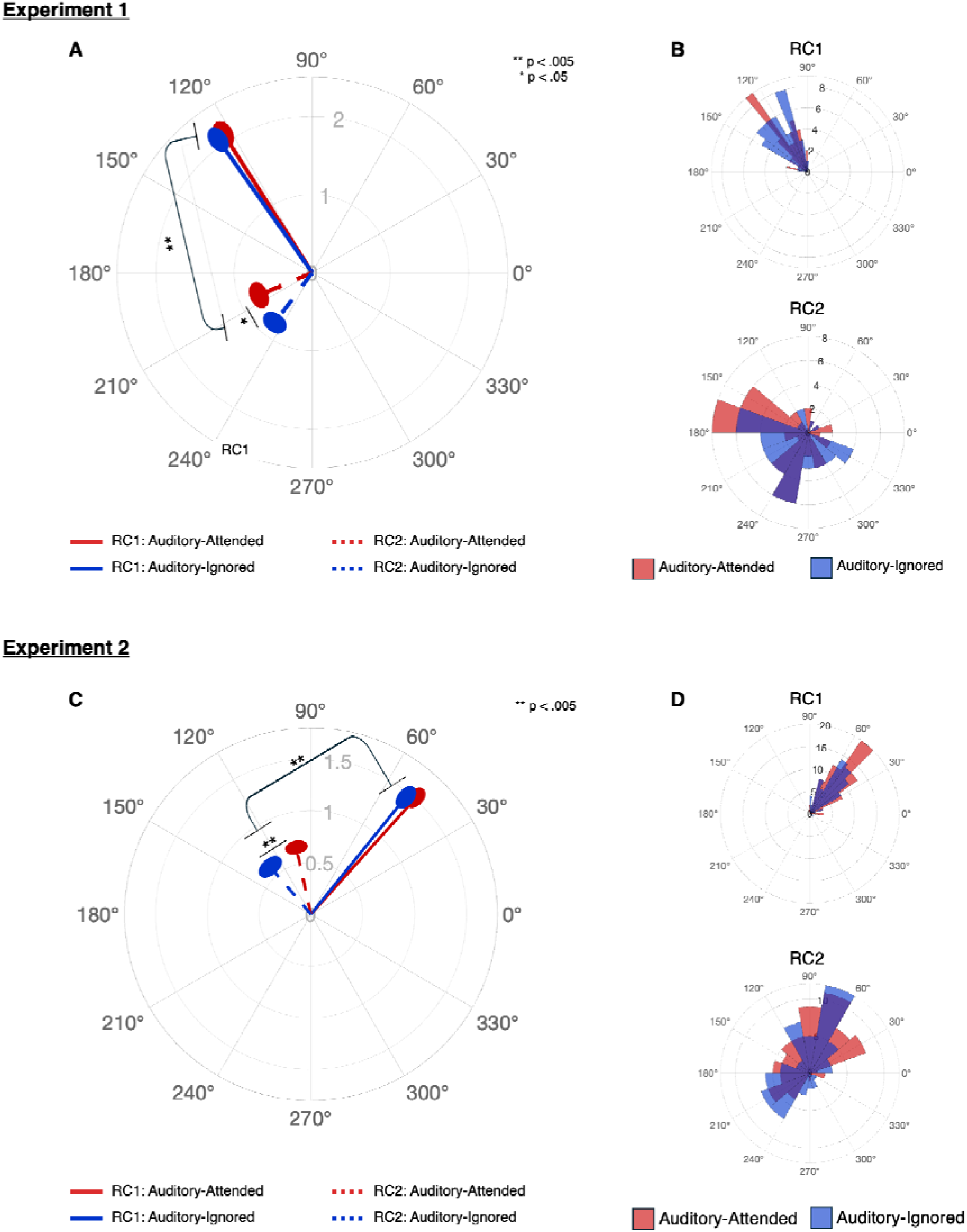
Phase alignment to the 3-Hz auditory stream across components and conditions. Top panel: Experiment 1; Bottom panel: Experiment 2 (including both sessions). Polar plots show the phase shift as an angle around the circle, with degrees increasing counterclockwise. The distance from the center represents amplitude, and the direction indicates the phase angle relative to a reference point (e.g., zero value). **Figures A and C** show for each experiment, group level circular phase-amplitude plot. Lines represent the phase alignment of the component-space data toward the 3-Hz auditory stream (RC1 as solid lines; RC2 as dashed lines) in each condition (‘Auditory-attended’-red;‘Auditory-ignored’ - blue). In both Experiments no significant condition-related phase differences were observed in RC1, whereas in RC2 there were consistently smaller 3-Hz phases in the auditory-attended condition. Ellipses indicate the standard error of the mean (SEM) for amplitude and phase. Asterisks denote significant differences between conditions. **Figures B and D** are polar histograms that show the distribution of the 3-Hz phase values of the component-space data (RC1 on the top; RC2 on the bottom) for both conditions. The radial axis indicates the cumulative number of participants contributing to each angular bin. Across all figures, red represents the Auditory-attended condition, and blue represents the Auditory-ignored condition. The zero reference point in the polar plots is arbitrary and should be interpreted only relative to the within-experiment contrasts.

### Individual-Level Temporal Alignment Is Stable Across Sessions

The test–retest design of **Experiment 2** enabled an assessment of the reliability of attention-related modulation of temporal dynamics across sessions. For each component, we computed the phase difference between the Auditory-attended and Auditory-ignored conditions and examined the stability of these differences across sessions using correlation analyses. Both components exhibited strong within-subject consistency, with high correlations observed for RC1 (*r* = 0.94, *p* <.001) and RC2 (*r* = 0.63, *p* <.001). Although the two components spanned different phase ranges, the slopes of the linear fits did not differ significantly (t_(34)_ = 0.8, p =.40), indicating comparable levels of test–retest consistency across components.

### Inter-Trial Phase Coherence Reflects Sensory-Driven Entrainment

In **Experiment 1**, a two-factor repeated-measures ANOVA (RC × Condition) revealed a significant main effect of RC (F_(1, 42)_ = 181.2, *p* <.001), indicating that ITPC was stronger in the RC1 component space compared to RC2. No other effects were observed.

In **Experiment 2**, a two-factor repeated-measures ANOVA (RC × Condition) also revealed a significant main effect of RC (F_(1, 71)_ = 131.55, *p* <.001), indicating stronger ITPC in RC1 compared to RC2. In contrast to Experiment 1, results also revealed a main effect of condition (F_(1, 71)_ = 11.43, *p* =.001), showing that ITPC was higher in the *Auditory-attended* condition compared to the *Auditory-ignored* condition. Across experiments, no significant interaction effects were detected. See Figure 3 for ITCP results.

**Figure 3:**
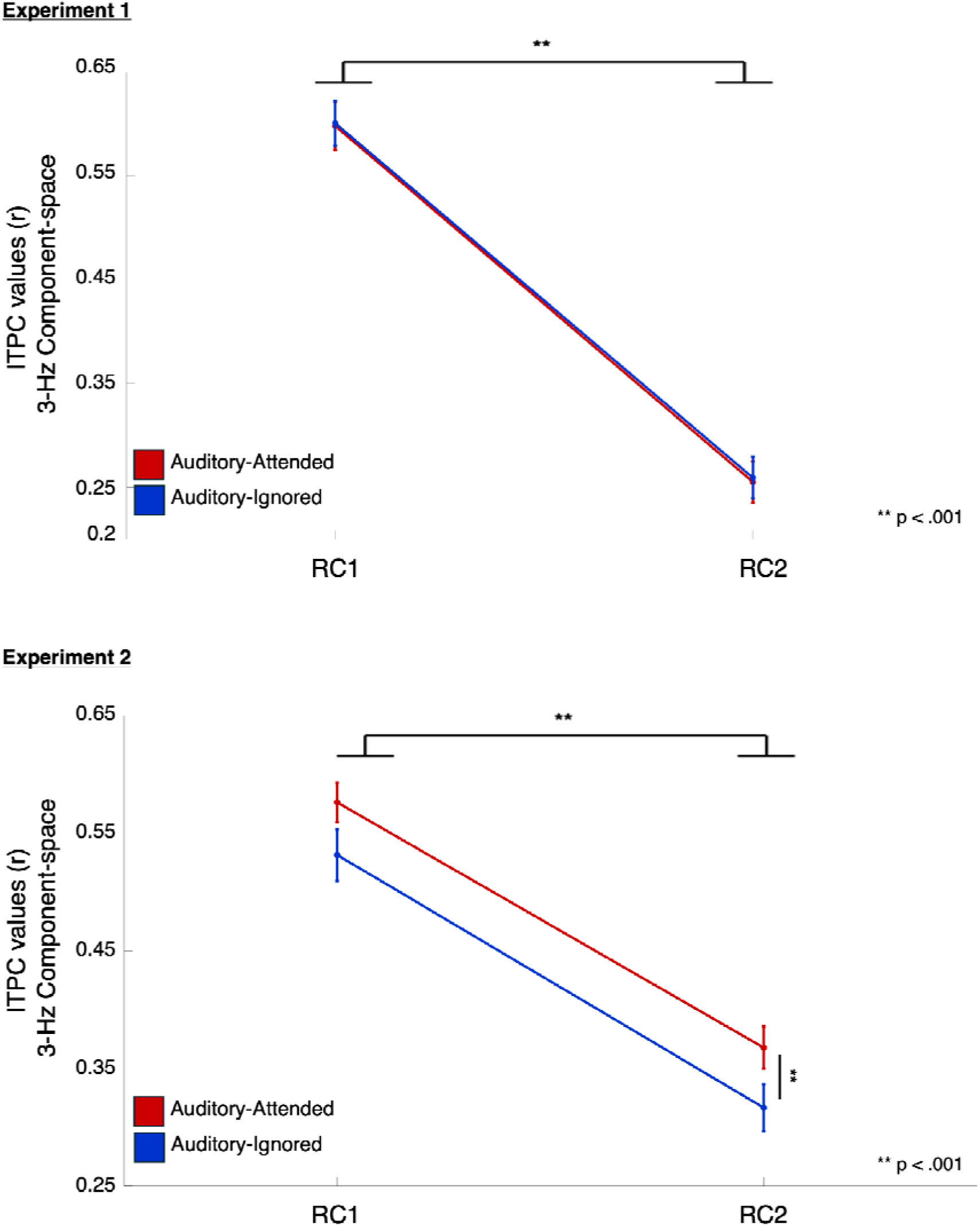
Inter-trial phase coherence (ITPC) at 3-Hz as a function of task condition and RC in both experiments. **Top panel:** Experiment 1; **Bottom panel:** Experiment 2. Line plots represent the ITPC across Auditory-attended (red) and Auditory-ignored (blue) conditions for each component. Bounding the line plots indicates the standard error of the mean (SEM) for ITPC. Asterisks denote significant differences between conditions and/or components.

### Individual Differences in Attentional Control Are Reflected in RC2 Temporal Dynamics

Our last step was to determine whether the different component levels we identified, and the distinct patterns of neural entrainment expressed within each, also relate to standardized measures of attentional ability. This analysis was only conducted with the participants of **Experiment 2**. Thus, we examined the relationship between the attentional modulation of neural entrainment to the auditory stream and participants’ performance on the NEPSY-II Auditory Attention subtest (AA-RS), a standardized neuropsychological assessment of auditory attention. Based on group-level averages, participants were categorized into high-and low-attentional-control groups. Circular ANOVA (RC × Group) was then conducted to test whether the 3-Hz component-space phase values (across the two sessions) differed between these groups.

Results revealed a main effect for group [*F*_(1, 69)_ = 4.24, *p* <.05] and an interaction effect [*F*_(1, 69)_ = 4.2, *p* <.05]. Post hoc paired-sample Hotelling’s tests showed that, relative to the low attentional-control group, individuals with higher attentional control exhibited smaller mean 3-Hz phase angles in the RC2 component space [*F*_(1, 32)_ = 4.83, *p* <.05], whereas no such group differences were observed in RC1 [*F*_(1, 32)_ = 0.1, *p* =.9]. These findings show that RC1 and RC2 capture distinct aspects of auditory processing, each showing a different pattern of correspondence with behavioral indicators of attentional modulation (see Figure 4).

**Figure 4:**
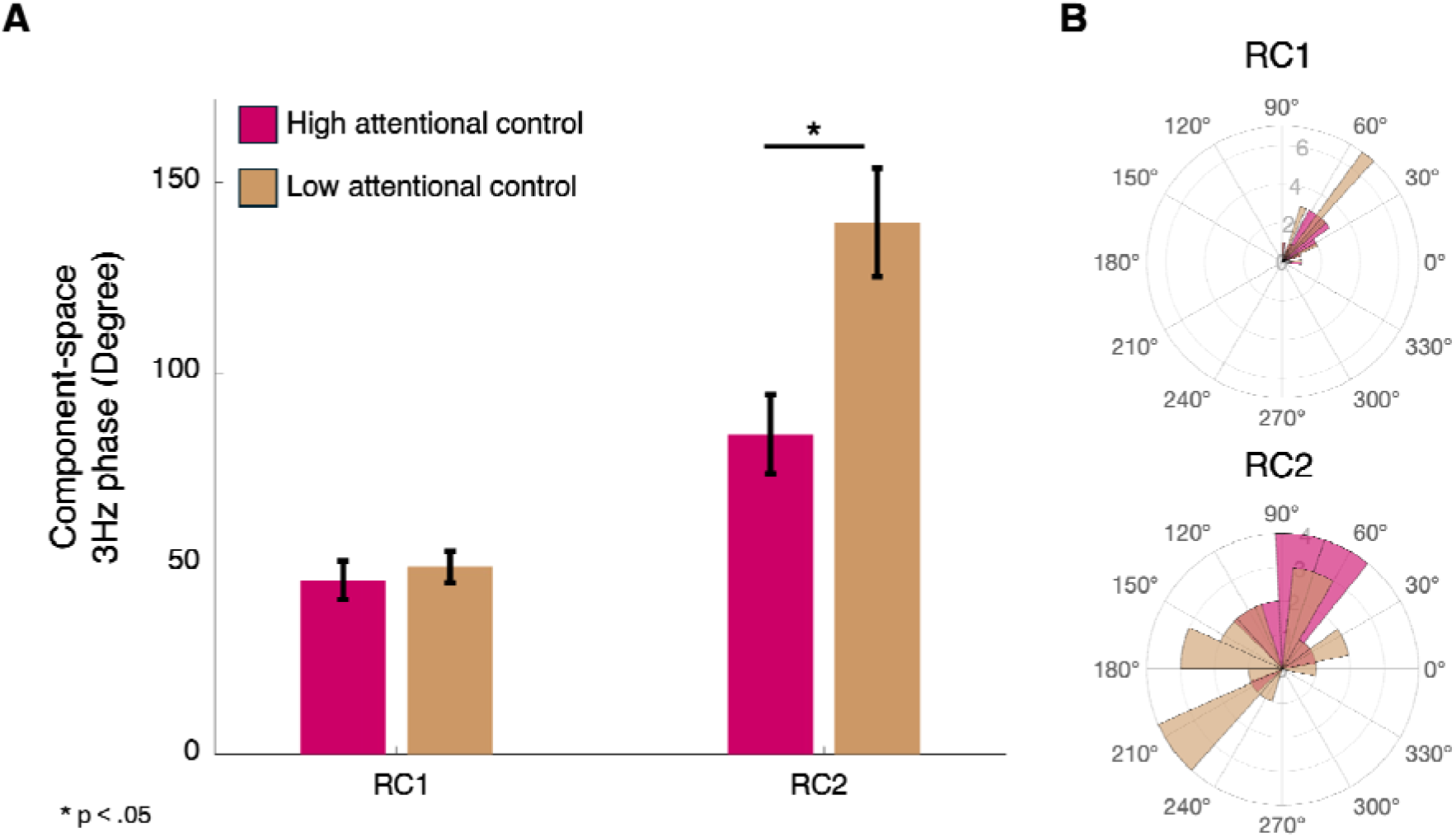
Group differences in 3-Hz phase shift as a function of attentional control. **(A)** Bar graph shows mean 3-Hz phase values (in degrees ± SEM) for RC1 and RC2, separated by attentional control level as measured through the Auditory Attention and Response Set (AA–RS) subtest of the NEPSY-II. Children with high attentional control (magenta) and low attentional control (tan) differed primarily in RC2, reflecting better phase alignment and shorter phases in the high-control group. Asterisks denote significant differences between conditions (p <.05).**(B)** Polar histograms show the distribution of the 3-Hz phase values of the component-space data (RC1 on the top; RC2 on the bottom) for both groups (high attentional control - magenta; low attentional control – tan). The radial axis indicates the cumulative number of participants contributing to each angular bin.

## Discussion

The current study advances a theoretical model suggesting that attention modulates auditory processing through Selective Temporal Alignment of Components (STAC). Across a series of highly replicable studies, we provide empirical evidence that attentional modulation of auditory processing is expressed selectively across dissociable hierarchical components. We identified two reliable components that differed in scalp topography, sensitivity to attention allocation, and behavioral relevance. The first component (RC1) exhibited stimulus-locked responses that closely tracked the rhythmic structure of the auditory input and showed minimal influence of attention. In contrast, a second component (RC2) displayed different temporal patterns of entrainment and robust attentional modulation: when attention was directed toward the auditory stream, 3-Hz phases systematically shifted toward smaller phase values, indicating a reorganization of the temporal alignment of neural activity with the auditory input. Consistent with models proposing that attentional control operates through higher-level mechanisms, we further show a relationship between individual differences in attentional control, as assessed by standardized measures, and attention-related shifts in temporal dynamics captured by RC2.

These results provide clear evidence for separability at the level of neural organization by showing that the attention-modulated network observed here (RC2) cannot be explained as a delayed or transformed version of the stimulus-driven component (RC1). On the contrary, RC2 reflects a distinct control process, suggesting that attention selectively targets higher-order networks modulating their temporal dynamics in response in a goal-directed fashion. Prior work examining attention-related changes in entrainment strength has shown that neural entrainment is not confined to primary sensory cortices but instead reflects the engagement of hierarchically organized systems spanning temporal, parietal, and frontal regions (Besle et al., 2011; Farahani et al., 2017; Kasten et al., 2024; Manting et al., 2020; Saupe et al., 2009). Although our component-space EEG analyses do not allow precise anatomical localization, the current findings extend this line of work by demonstrating that hierarchical organization is also evident in the temporal domain, with attention systematically reorganizing the timing of neural entrainment rather than merely amplifying its magnitude. In addition, our results indicate that these functionally distinct networks operate on different temporal scales: the automatic, stimulus-driven response is rapid and robust, yielding a strongly similar response across participants, whereas the attention-related component also entrains to the auditory stimulus but unfolds more slowly and shows greater inter-individual variability. This dissociation suggests attentional modulation acts as a process that flexibly adjusts the timing of neural activity according to task demands and individual attentional strategies, rather than uniformly amplifying a shared sensory response.

Importantly, our results show that temporal realignment is not a byproduct of attentional engagement, but a core operation through which attentional control is implemented. Single-cell recording studies in animal models have shown that attentional control in multisensory environments operates by adjusting the temporal timescale of oscillatory activity within higher-order processing systems, while leaving early sensory processing largely stimulus-driven (O’Connell et al., 2014; Schroeder & Lakatos, 2009; Zuo et al., 2020). Our findings echo these insights, highlighting the role of phase shifts and temporal reorganization as a central mechanism through which attention can shape cognitive processing. In human studies, this temporal dimension has received relatively little attention, as the majority of work has relied on phase-consistency measures such as inter-trial phase coherence (ITPC) that primarily index the reliability of entrainment within single trials. These studies have shown that directing attention toward repetitive auditory stimuli increases ITPC, reflecting more reliable stimulus-locked neural responses (Guerra et al., 2024; Kachlicka et al., 2022; Laffere et al., 2020, 2021). However, these measures do not provide information about the phase position at which entrainment occurs relative to the stimulus, limiting their ability to capture directionally specific, attention-driven changes in temporal dynamics. STAC moves beyond measures that are insensitive to systematic shifts in phase angle by dissociating neural responses into components and directly assessing dynamic shifts in phase alignment within each. In doing so, it provides a mechanistic account of how attention modulates sensory processing, rather than merely increasing the reliability of stimulus locking.

From this perspective, ITPC and STAC capture fundamentally different aspects of how attention modulates neural tracking. ITPC captures the stability of stimulus-driven phase alignment across repetitions, whereas STAC characterizes whether attentional engagement induces systematic, directionally specific shifts in phase timing. This distinction is directly illustrated by our results showing that ITPC effects peaked in the stimulus-driven component (RC1), consistent with reliable sensory synchronization to rhythmic input, whereas attention-related phase shifts were most pronounced in the attention-modulated component (RC2), reflecting top-down adjustments in oscillatory timing. In addition, while ITPC-based attentional effects varied across experiments, the phase-shift measures derived from STAC were robust both within and across individuals, underscoring the value of explicitly modeling temporal reorganization when studying attentional mechanisms.

One important implication of the STAC framework is its potential to serve as a stepping stone for advancing theories of temporal prediction, providing a fine-grained method for explicitly examining how attention shifts neural entrainment to dynamically prioritize specific moments or units within the auditory and linguistic stream (Lakatos et al., 2019; Park et al., 2015). Previous work in animal models has shown that attention can realign oscillatory phase to expected sound onsets and thus enhance temporal prediction and perceptual sensitivity (Henry & Obleser, 2012; Lakatos et al., 2008; Zoefel & VanRullen, 2016). A growing body of research suggests that similar mechanisms operate during natural speech processing, with neural entrainment to speech rhythms supporting segmentation, prediction, and comprehension across multiple linguistic levels (Ding & Simon, 2014; Giraud & Poeppel, 2012; Peelle & Davis, 2012).

In the present study, we employed steady-state auditory stimuli composed of syllables presented at a speech-like rhythmic rate (∼3-Hz) in a free-field setting. Although the stimuli were composed of discrete syllables, they preserved core temporal and acoustic features of speech, including rhythmic structure and syllabic rate, allowing us to examine how attention modulates the timing of neural phase alignment in response to speech-like input. We demonstrate a consistent phase shift of approximately 50° when attention was directed toward the stimuli, across two independent cohorts. In the current design, syllables were presented in a 3-Hz stream, with each syllable lasting approximately 200 ms and followed by a short silent gap. Within this temporal structure, the observed ∼50° phase shift corresponds to an advancement of roughly 45 to 50 ms in neural timing, suggesting that attention promotes an anticipatory alignment of oscillatory phase just before the onset of the upcoming syllable. In the context of the present task, in which participants were required to detect targets composed of two consecutive syllables, such temporal realignment is well-positioned to facilitate the integration of successive syllabic information, consistent with a predictive timing mechanism rather than a purely reactive sensory response. Although the current paradigm employs simplified, syllable-based stimuli, it preserves the essential rhythmic and phonological dynamics of speech while minimizing linguistic processing demands. This design is therefore well-suited for isolating how attentional and temporal mechanisms contribute to temporal prediction, without confounding influences from more complex linguistic or cognitive factors. Leveraging STAC, we further show that attentional modulation of neural phase timing extends beyond stimulus-locked synchronization to support predictive alignment processes that are critical for efficient, goal-directed speech parsing.

Finally, the present findings extend the study of attentional and temporal dynamics into a developmental context by examining these mechanisms during adolescence, a period in which sensory regions supporting auditory processing are largely mature, whereas frontal and frontoparietal networks implicated in attentional control continue to undergo substantial refinement (Calcus, 2024; Karns et al., 2015; Lackner et al., 2013; Lalonde et al., 2024). Our results show attention-related temporal effects across age groups, suggesting that our model is sensitive to attentional shifts in temporal dynamics in different developmental stages.

This is particularly important given that children and adolescents with attentional difficulties frequently struggle to focus on target speech in the presence of competing auditory input (Blomberg et al., 2019; Dawes & Bishop, 2009; Laffere et al., 2021; Lemel et al., 2023; Levy et al., 2025; Michalek et al., 2014). However, it remains unclear whether these difficulties stem from core impairments in attentional control or from the additional linguistic and cognitive demands imposed by complex listening environments. By employing speech-like rhythmic stimulation with minimal linguistic load, the present paradigm helps disentangle these possibilities and offers a sensitive framework for probing the development of temporal attentional mechanisms.

Importantly, STAC yields robust and highly replicable effects within an exceptionally brief paradigm (less than two minutes in total), making it particularly well-suited for research with young children and populations with neurodevelopmental disorders, where sustained engagement can be challenging. While the current study focuses on adolescents, future work directly comparing age groups within a unified design will be essential for clarifying how these temporally organized attentional processes evolve across development and how individual differences relate to real-world listening and learning outcomes.

### Limitation and Future directions

Several limitations should be noted. First, although sensor-level RCA allows us to dissociate functionally distinct components, it does not permit precise anatomical localization of the underlying neural sources. Thus, conclusions regarding hierarchical organization are based on functional dissociation rather than direct source reconstruction. Second, the current sample focused on adolescents, limiting strong developmental inferences across broader age ranges. Direct cross-age comparisons within a unified design will be necessary to more precisely characterize developmental trajectories. Finally, while the paradigm was intentionally minimal and brief to maximize signal-to-noise ratio and feasibility, this controlled design may not fully capture the complexity of real-world listening environments. Future work integrating STAC with more naturalistic stimuli will help bridge this gap.

## Conclusions

The current study identifies temporal reorganization as a central mechanism through which attention shapes auditory processing. Using a new theoretical and methodological framework, Selective Temporal Alignment of Components (STAC), we show that attention modulates neural entrainment not only by strengthening stimulus-locked synchronization or modulating entrainment strength, but by selectively shifting the timing of higher-order oscillatory activity in anticipation of behaviorally relevant information. This finding reframes attentional modulation in terms of *when* neural activity is aligned, rather than solely *how strongly* it is synchronized.

Our results further support a hierarchical model of attentional control, in which stimulus-driven entrainment and goal-directed modulation are functionally dissociable processes operating at distinct temporal scales. By resolving neural activity into components, STAC reveals that attentional influences emerge primarily through temporally structured adjustments in higher-order networks, while lower-level sensory tracking remains tightly locked to the stimulus. This component-level separation clarifies how top-down and bottom-up processes interact during rhythmic auditory processing, extending hierarchical accounts of attention into the temporal domain.

Finally, by examining these mechanisms in adolescence, the present work informs theories of speech processing and development. Neural entrainment to speech rhythms has been implicated in segmentation, prediction, and comprehension, yet how attentional control over temporal dynamics matures remains poorly understood. The present findings demonstrate that temporally organized attentional modulation is already evident during this developmental period, providing a framework for future work on how predictive timing mechanisms support the development of efficient speech processing and how their disruption may contribute to attentional and auditory processing difficulties.

## Methods

### Participants

#### Experiment 1

Forty-seven students, aged 10–12 years, participated in *Experiment 1*. Each participant completed a single EEG session in a school-based EEG lab (Toomarian et al., 2024). Participants were recruited from the school, and sessions were conducted during school days. All students whose parents provided written informed consent were allowed to participate in the study regardless of psychological or medical diagnoses; students also assented to participating. Demographic data and information on participants’ diagnosis, medical history, and current prescribed stimulant medication were collected from the parents and are reported in Table S1.

From this data set, three participants were excluded due to excessive motor artifacts (over 60% of the session), and one data set was removed following a technical issue during the session. Thus, the presented analysis shows data from 43 participants in total (age range: 10–12 years, M = 10.8, SD = 0.72; 22 girls).

#### Experiment 2

Forty-three students aged 12–14 years participated in this study, which included two sessions, 3-4 weeks apart, in the same school-based EEG lab as Experiment 1. Similar to the first experiment, participants were recruited from the school, and sessions were conducted during school days. All students with signed consent forms from their parents, and who provided assent, were allowed to participate (see Table 1 for demographic data and information regarding diagnosis, medical history, and current prescribed medication).

Five participants were excluded from the final data set: three presented excessive motor artifacts (over 60% of the session, across all trials combined), and two did not complete the full task design. Two additional participants’ data sets were removed due to technical issues that emerged in the first session. Thus, the presented analysis shows data from 36 participants in total (age range: 12–14 years, M = 12.9, SD = 0.6; 13 girls).

Both studies were approved by the Stanford University Institutional Review Board, and each child received a small token gift as compensation for participation.

### Stimuli

The current work consisted of two experiments using a shared audiovisual selective attention paradigm, with age-appropriate adaptations introduced as needed. We first describe the general task structure common to both experiments, followed by experiment-specific details.

In both experiments, participants completed an audiovisual selective attention paradigm that incorporated core elements of the continuous performance task (CPT; Riccio et al., 2002; Rosvold et al., 1956). Each trial consisted of a 48-second audiovisual movie featuring two concurrently presented rhythmic streams: a 3-Hz auditory stream composed of spoken syllables and a 1.25-Hz visual stream composed of letters appearing on a computer screen. Each auditory syllable was presented for approximately 110 ms in duration, with silent intervals inserted to establish a 3-Hz presentation rate. Visual stimuli appeared on the screen for 0.8 seconds to create the 1.25-Hz steady-state stream. The letters on the screen were presented without an intervening fixation cross between them to maintain the steady-state effects (De Rosa et al., 2022; Norcia et al., 2015; Wang et al., 2023).

The auditory stream was designed to approximate the temporal characteristics of natural linguistic input (Laffere et al., 2021; Poeppel & Assaneo, 2020). By constraining the auditory rate to 3-Hz, we emphasized syllable-level processing, which is ecologically relevant for speech perception in children. To avoid confounding the task with frequency-related processing difficulty, we followed previous studies (Dhooge & Hartsuiker, 2010; Miozzo & Caramazza, 2003) and selected lower-frequency presentation rates for both modalities. The specific frequencies of stimuli were chosen to ensure that each trial could be divided into twelve 4-second segments, with each segment capturing full cycles of both the 3-Hz auditory and 1.25-Hz visual streams, facilitating frequency-specific analyses within each segment (see data analysis).

In each trial, participants were asked to perform one of two tasks. In the *Auditory-attended* condition, participants were instructed to pay attention to the auditory syllables and press a button when they detected specific target pairs within it (e.g., ‘Ba’ followed by ‘Bo’). In contrast, in the *Auditory-ignored* condition, they were asked to shift their attention away from the auditory stream and instead identify specific targets in the visual stream, as described below. Audio and visual target pairs were separated by at least two other randomly chosen stimuli to reduce the likelihood of attentional blink effects (Shapiro et al., 1997). In addition to the target pairs, two types of distractor sequences were included in both the auditory and the visual streams, each comprising 12% of the stream: misleading initial distractors (e.g., ‘Bo’ followed by ‘Ga’/‘Go’) and misleading follow-up distractors (e.g., ‘Ga’/‘Go’ followed by ‘Ba’).

Although the overall paradigm was similar across both experiments, several task elements were adapted to suit the different age groups and practical constraints related to the main speaker (for reasons unrelated to the scope of this study). Thus, while comparable in structure, the tasks were not identical and should not be directly compared. In **Experiment 1,** the auditory and the visual streams comprised four consonant–vowel syllables - ‘Ba’, ‘Bo’, ‘Ga’, and ‘Go’- presented in a pseudo-random sequence with 14 targets in each stream (approximately 20% of auditory items; 45% of visual items). In both the *Auditory-attended* and the *Auditory-ignored* conditions, participants were instructed to detect the same target pair (‘Bo’ followed by ‘Ba’) in either the auditory or visual modality. In **Experiment 2,** we distinguished the two streams by changing the stimuli in the visual stream from syllables to four letters - ‘M’, ‘X’, ‘Z’, and ‘L’ in the *Auditory-attended* condition and ‘S’, ‘B’, ‘Q’, and ‘R’ in the *Auditory-ignored* condition. This was conducted to separate the auditory and the visual conditions from each other and prevent mixed signals between the streams. In addition, to maintain high levels of attention within the older-age group, we controlled the rates of the targets within each stream and raised them to 50% of the targets in each stream.

In both experiments, the auditory syllables were recorded by the students’ teachers: in the first experiment, the stimuli were narrated by the female lead teacher of the classes, and in the second by a male lead teacher of the older students’ classes. All stimuli were recorded within the school-based lab using a professional microphone and were reedited with Audacity (version 3.7; https://www.audacityteam.org/) to make sure the volume and the onset were equivalent between the different syllables. Custom-made MATLAB scripts were used to create the auditory and the visual streams, based on the criteria mentioned above, and all stimuli were presented using the Psychtoolbox package, MATLAB (R2024b, version 24.2.0).

### Experimental Procedure

In each session, participants were seated in a sound-attenuated room with lighting calibrated to be consistent across participants, at a viewing distance of approximately 1 m from a computer screen. The participants were informed of the current study goals and were familiarized with the EEG net. They were then asked to sign an assent form, in addition to the written informed consent that their parents provided before the study, to make sure they wanted to take part in this session.

After signing, students were told that their teacher needed their help, and their goal was to assist by identifying target syllables in the auditory or visual stream (*‘Auditory-attended’* and *‘Auditory-ignored’* conditions, respectively). They were then trained on both conditions to ensure that the instructions were clear.

Each trial started with a 2-second on-screen instruction cue indicating the upcoming task (*Auditory-attended* or *Auditory-ignored*), followed by a 4-second short training period that contained ramping up unimodal stimuli - auditory or visual, based on the condition for the current trial. The trial then proceeded with the 48-second presentation of the main multimodal stimuli (see Figure 5B). All visual stimuli were presented at the center of the screen in a white font on a gray background. Auditory stimuli were played in a free-field manner through a speaker that was located in front of the participant, above the screen. Participants were asked to press a button on an external response pad with their preferred hand when they identified the target (see Figure 5C). During both conditions, participants were asked to keep their eyes on the screen, and their gaze direction was monitored by a research team member to ensure that they were keeping their gaze toward the center of the screen. Participants were given verbal feedback about their performance and the consistency of their gaze after the end of each trial. Overall, participants completed six trials in Experiment 1 and eight trials in Experiment 2, with trials split evenly across the two attention conditions. Trials were pseudo-randomized across participants and organized into separate blocks within each session. Each block contained one *Auditory-attended* trial and one *Auditory-ignored* trial presented in random order (see Figure 5A). In both experiments, between the blocks, participants engaged in a narrative-listening activity, which is beyond the scope of the present paper. EEG sessions took approximately 55 minutes, including setup and between-trial breaks.

**Figure 5:**
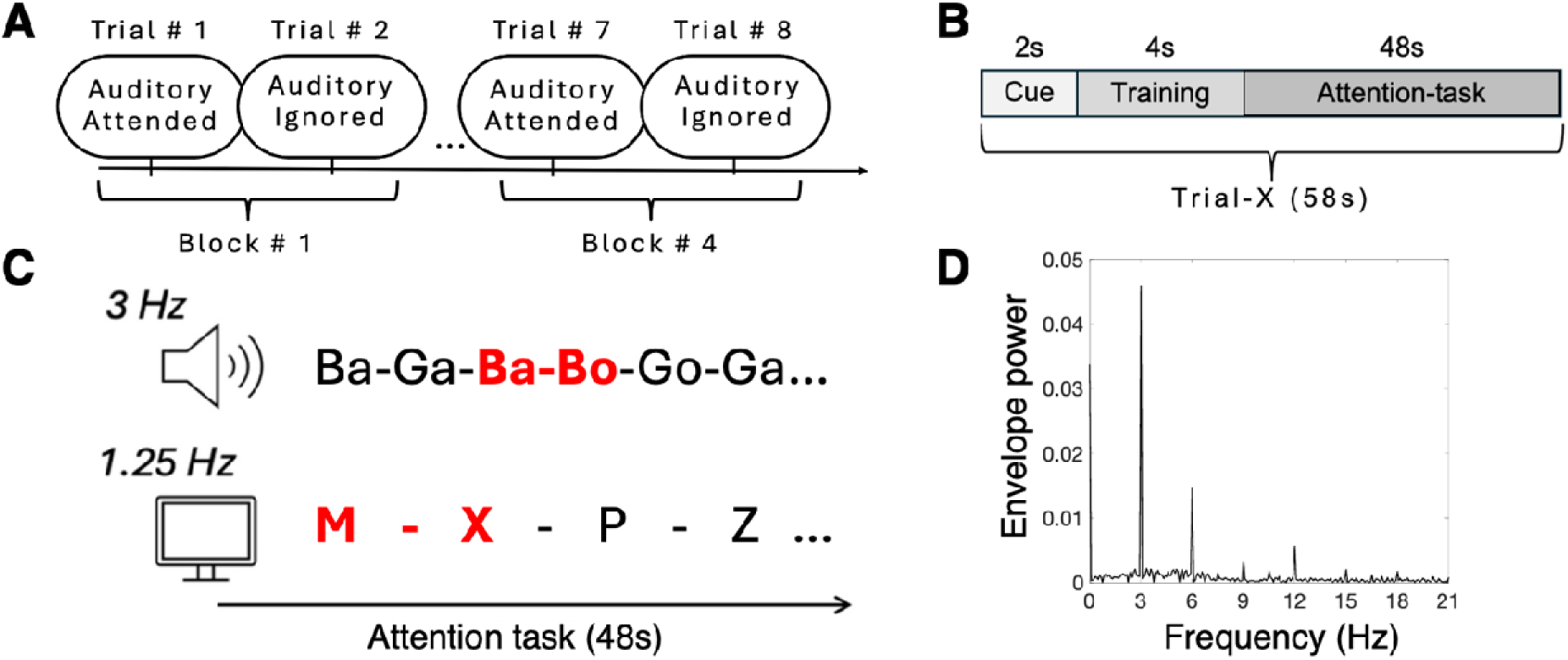
Experimental procedure. **A.** Schematic illustration of one experimental session. Both experiments consisted of separate blocks with short breaks in between. Within each block, participants completed two trials: one auditory-attended trial, in which they were instructed to focus on the auditory stream, and one auditory-ignored trial, in which they were instructed to ignore the auditory input and attend to the visual stimuli presented on the screen. **B.** Schematic illustration of a single trial, beginning with a cue indicating the condition (auditory-attended or auditory-ignored), followed by a short training period in which only the target stream was presented (auditory-only or visual-only). The main part of the session was 48 seconds, including both the auditory and the visual stimuli **C.** The attention task: a 48-second movie composed of 3-Hz auditory stimuli and 1.25-Hz visual stimuli, presented simultaneously. Participants were asked to identify specific targets (examples highlighted in red) using a button press. **D.** Sound envelope spectra of an example movie, showing clear peaks at 3-Hz and its harmonics.

### Cognitive assessment

In addition to the EEG sessions, in **Experiment 2,** participants completed a separate assessment session that included a battery of cognitive and reading assessments, as part of a broader effort to collect standardized data for each student. For the current study, we used scores from the Auditory Selective Attention subtest of the NEPSY–II. This well-established standardized test for children and adolescents evaluates the ability to control auditory attention in a multisensory audiovisual environment through two components. Part A (*Auditory Attention; AA*) requires students to sustain attention while listening to a list of words and to press a colored circle in response to a specific target word (e.g., *“press the red circle when you hear the word red”*). Part B (*Response Set; RS*) adds higher demands on executive function and auditory control, as children must inhibit the previously learned response and apply a new, more complex rule (e.g., *“press the red circle when you hear the word yellow”*). For analysis, we used the combined standardized AA-RS score, which integrates performance across both parts to provide an overall measure of auditory attentional control.

### EEG Recording

Neural activity was recorded using a 128-sensor EEG net (Electrical Geodesics, EGI NA 400 amplifier) against the Cz reference, and was sampled at 1000 Hz. Electrooculographic (EOG) signals were measured by 4 facial electrodes located above and on the external side of both eyes. Impedances were kept below 50 kΩ (Ferree et al., 2001). Data were then imported into custom-made MATLAB codes for preprocessing.

## Data Analysis

Data analysis was identical for the two experiments, unless explicitly mentioned otherwise.

### Behavioral Data Analysis

To evaluate the overall performance in the task, we calculated the accuracy and the reaction times toward targets across all task blocks for each one of the task conditions. To assess target detection, we analyzed responses within a 700-ms window following the presentation of the second stimulus in each pair. A response was considered correct if it occurred within this window; responses outside of this window or responses made earlier than 50 ms after the window began were classified as false alarms. High levels of accuracy rates were interpreted as evidence of sustained attentional engagement, while increased error rates suggested potential attentional lapses. In **Experiment 1**, paired-samples t-tests were used to compare accuracy and reaction times between conditions. In **Experiment 2**, since participants completed two sessions, we used repeated-measures ANOVA (Condition × Session) to examine performance while explicitly accounting for session-related differences.

### EEG Data Analysis

#### Preprocessing

EEG data were preprocessed using the FieldTrip toolbox (MATLAB, https://www.fieldtriptoolbox.org, version 20220729). For each participant, the raw EEG data from all trials (both conditions) were appended together for the preprocessing phase. First, all data were band-pass filtered between 0.5 and 40 Hz (4th-order zero-phase Butterworth IIR filter), detrended, and demeaned. Visual inspection was performed to identify and remove gross muscle artifacts, and Independent Component Analysis (ICA) was then used to further remove components associated with horizontal or vertical eye movements and heartbeats. Remaining noisy electrodes, containing extensive high-frequency activity or DC drifts, were removed and interpolated using neighboring electrodes.

After artifact rejection, data were resampled to 420 Hz to ensure an integer number of frames per stimulation cycle, corresponding to the auditory and the visual stimuli frequencies of 3-Hz and 1.25-Hz (i.e., 20 frames for 3-Hz, 48 frames for 1.25-Hz). Following this, the continuous EEG recordings were re-referenced to an average reference configuration (Lehmann & Skrandies, 1980) and segmented into 4-second epochs, resulting in 12 segments per block. The first and the last epochs of each trial were removed, and noisy epochs were excluded based on visual inspection. These data are henceforth referred to as the *broadband* data.

Finally, to examine neural entrainment to the auditory stimuli solely, we filtered the data to retain activity from only the first five harmonics of the 3-Hz target auditory frequency (3, 6, 9, 12, and 15 Hz). We applied a Fast Fourier Transform (FFT) to convert the signal from the time domain to the frequency domain, zeroed out the values at all positive and negative frequency bins representing non-target frequencies, and then performed an inverse FFT to return the filtered data to the time domain for subsequent analyses. Thus, the time-domain *frequency-filtered* data are matched in size to the broadband data, although activity from only the target frequencies is represented in the frequency-filtered data.

#### Reliable components analysis (RCA)

The first step of the STAC analytic approach aimed to identify reliable spatial components that reflect distinct levels within the hierarchical organization of auditory neural entrainment. We applied Reliable Components Analysis (RCA) to the 3-Hz *filtered data*. RCA is a data-driven spatial filtering approach that extracts neural components by maximizing covariance across multiple data records (e.g., trials or participants). By transforming the full 128-channel sensor space into a smaller set of weighted components, this method achieves a higher signal-to-noise ratio (SNR) compared to other approaches and enables the identification of consistent response patterns without requiring manual channel selection (Dmochowski et al., 2012, 2015). In the current study, we used RCA not only to identify distinct neural components corresponding to auditory entrainment but also to use these components as spatial filters to examine how attention modulates the temporal dynamics within distinct neural networks. To identify the neural components that represent auditory processing independent of attentional allocation, RCA was performed on the filtered data from both conditions combined. The analysis was conducted using a publicly available MATLAB implementation^1^ (Dmochowski et al., 2015). Five reliable components (RCs) were extracted, with regularization applied by retaining the first seven principal components from the pooled within-trials covariance eigenvalue spectrum. The computation results in a sensor-by-component weight matrix *W* (i.e., the eigenvectors), along with a vector *dGen* of associated eigenvalues representing trial-to-trial correlations. These eigenvector-eigenvalue pairs are ordered in descending order of ‘reliability’ explained – i.e., RC1 is the maximally correlated component. To generate scalp topographies of the spatial filters, we computed the forward-model projection matrix *A* from the calculated *W* and the pooled within-trial covariance matrix, following methods from Parra et al. (2005).

To guide RC selection, we assessed the statistical significance of each component’s eigenvalue using permutation testing. A null distribution was generated by repeating the RCA calculations on phase-scrambled EEG data, a procedure that disrupts temporal structure while preserving autocorrelation and power spectra (Kaneshiro et al., 2020; Prichard & Theiler, 1994; Wang et al., 2023). We generated null distributions by applying random phase rotations (0 to 2π) to the real and imaginary coefficients at each selected frequency, across all electrodes and trials, and then computing RCA; this was done 500 times, with phase rotations independently randomized each time. Observed eigenvalues were compared against the null distributions to obtain p-values, with multiple comparisons controlled using False Discovery Rate (FDR) correction across five tests (Benjamini & Hochberg, 1995). Based on this analysis and the observed topographic patterns, in both experiments, RC1 and RC2 were selected for subsequent analyses. These components reflect spatial patterns across conditions that capture maximal trial-to-trial covariance in phase-locked neural responses to the auditory stimulus frequency.

Next, to characterize the effects of auditory attention on the temporal dynamics within each RC, we applied the group-level RC1 and RC2 as spatial filters (128 weights) to each participant’s broadband sensor-space EEG data. The broadband preprocessed time-domain data from each condition (*Auditory-attended* and *Auditory-ignored; N* time points * 128 channels) were projected through these filters, yielding four component-space time series: *RC1-Auditory attended; RC2-Auditory attended; RC1-Auditory ignored; RC2-Auditory ignored*. (For full analytic pipeline see Figure 6)

**Figure 6:**
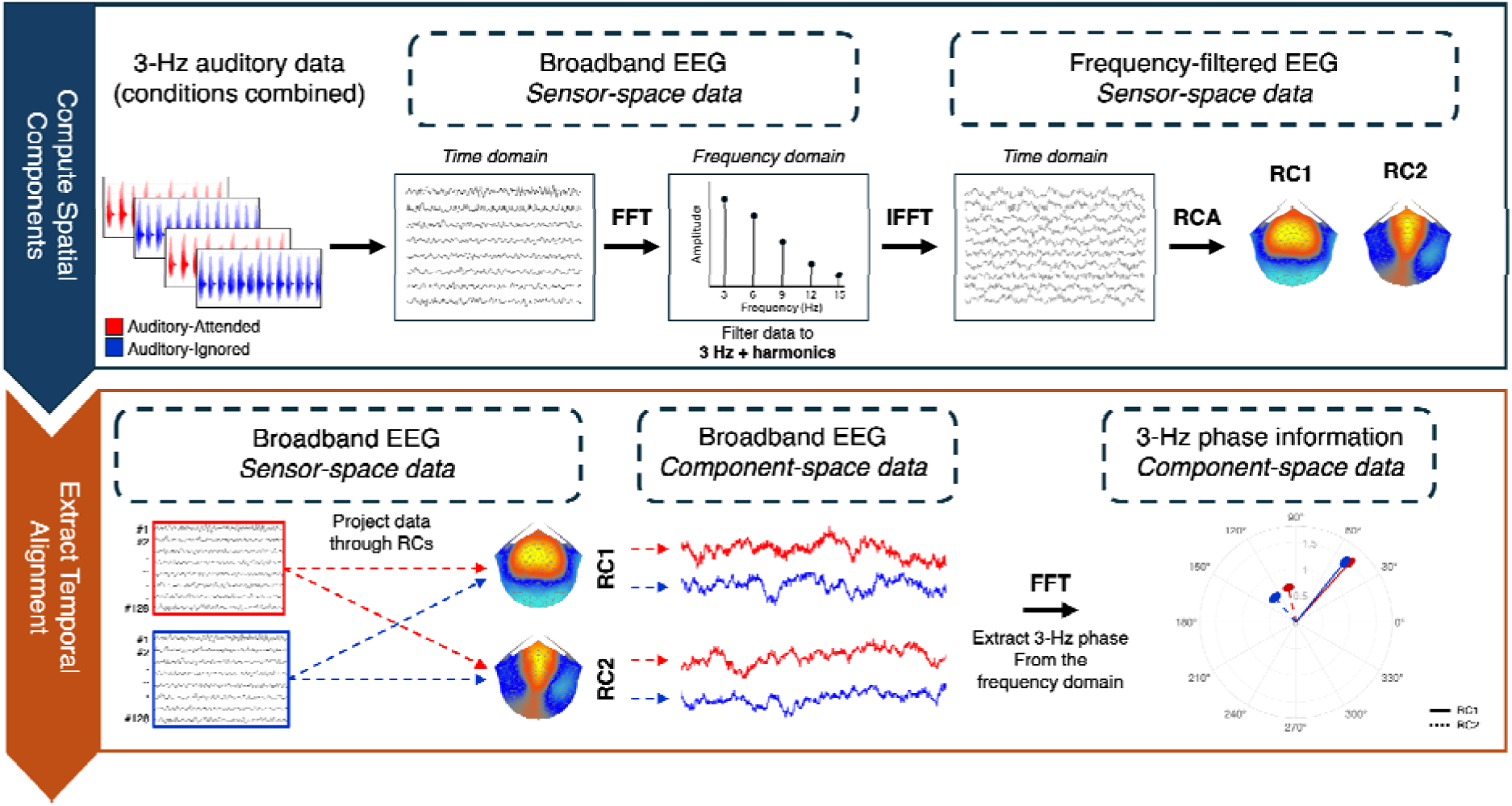
Overview of the analytic pipeline. **Top panel (Compute Spatial Components):** Broadband EEG data from the Auditory-Attended and Auditory-Ignored conditions were combined and transformed to the frequency domain using FFT. The frequency-domain data were then filtered to the stimulation frequency (3 Hz) and its harmonics (6, 9, 12, 15 Hz), followed by inverse transformation (IFFT). Finally, RCA was then applied to identify spatial components (RC1, RC2) capturing consistent 3-Hz auditory responses independent of attentional allocation. **Bottom panel (Extract Temporal Alignment):** For each condition separately, broadband sensor-space EEG was projected through the significant RCs to obtain component-space signals. These component-domain data were transformed to the frequency domain (using FFT), and the phase of the 3-Hz response was extracted. Circular statistics were used to compare phase distributions between attentional conditions, quantifying attention-related temporal dynamics effects.

### Assessing attention-driven modulation within components

To explore whether neural temporal dynamics are modulated by auditory attention, we explored the 3-Hz phase in each of the four component-space time series. FFT analysis was applied to each participant’s component-space data, and we extracted the phase and amplitude at the 3-Hz frequency. Circular statistics were used to assess attention-related temporal modulation across components, with a paired-sample Hotelling test evaluating whether the 3-Hz phase differed across task conditions and RCs. To measure RC × Condition interaction effects, we conducted a paired-sample Hotelling test applied to the difference-of-differences contrast (RC1−RC2) − (*Auditory-attended* − *Auditory-ignored*). Analyses of phase differences included both phase and amplitude measures to ensure that any observed effects were not driven solely by amplitude variations. Due to multiple paired-sample tests, all resulting p-values were corrected using a Bonferroni correction. All statistical analyses and computations were performed using circular statistics toolboxes (Mathworks, https://www.mathworks.com; Berens, CircStat, 2009; Rudy van den Brink, circ_htest, 2014) and custom-made MATLAB scripts. We also assessed whether the 3-Hz amplitude significantly differed across task conditions and RCs by performing a 2×2 (RC × Condition) repeated-measures ANOVA on the amplitude values at 3-Hz. The full analysis and complementary figure are provided in the Supplementary Information.

### Experiment 2: Individual-level consistency of attention modulation across sessions

Experiment 2 was designed to probe the consistency and individual interpretation of attention-related temporal dynamics using a test-retest design and cognitive measures. Accordingly, in this experiment, we examined individual-level consistency across sessions.

To evaluate the consistency of attention-related effects on neural temporal dynamics, we examined the test-retest reliability across the two EEG sessions in Experiment 2. Specifically, we assessed whether individuals’ ability to shift attention and more effectively track target auditory stimuli remained stable across sessions. For each component, attention-related modulation was quantified as the difference in 3-Hz phase values between the *Auditory-attended* and *Auditory-ignored* conditions. We then conducted circular correlations on these phase differences across the two sessions, separately for each component.

In addition to assessing the temporal dynamics in each component, we also explored whether inter-trial phase coherence (ITPC) differs between components and conditions. Previous studies (Kachlicka et al., 2022; Laffere et al., 2020, 2021) have shown that auditory attention increases the temporal alignment of neural oscillations across trials. This measure, which highlights the consistency and precision of neural phase alignment over time, provides a complementary perspective to our analyses by showing how consistent the neural phase alignment is over time in each processing level. Similar to the evaluation of the impact of attention on the temporal dynamics, ITCP was computed on the component-space data for each component and condition separately. We used a 4-second sliding window to segment the data. The window size was based on the auditory and visual frequencies, allowing an integer number of cycles for each. Then, a Hann-windowed FFT was applied to each segment, resulting in a complex vector at 3-Hz that was converted to a unit vector. All unit vectors from all segments were averaged, and the length of the resulting mean vector was taken as the ITPC value. This measure ranges from 0 to 1 (from no phase consistency across trials to perfect phase consistency, respectively). To assess attentional modulation of ITPC, we conducted a two-way ANOVA with Condition and Component as within-subject factors.

### Standardized Measures of Auditory Attention

In further support of Experiment 2’s aims toward individual interpretations and better understanding how our results, which focused on steady-state stimulation, relate to individuals’ ability to allocate auditory attention across contexts, we also examined how our neural measures mapped onto standardized neuropsychological assessments. To that end, we used the *Auditory Attention* and *Response Set* (AA–RS) subtest from the NEPSY-II battery (Korkman et al., 2012). Similar to the task we used in the EEG sessions, this assessment requires participants to sustain auditory attention over time, monitor auditory input while processing visual stimuli, and adjust to changing rules. Cognitive assessments were conducted in a separate session, and participants who did not complete this session were excluded from the analysis (n = 2).

Based on participants’ standardized scaled scores, we divided the sample into two subgroups to characterize distinct attention styles: participants with high sustained attention and cognitive control abilities, and participants who have difficulties maintaining their auditory attention in a multisensory environment. For this comparison, we focused exclusively on the *Auditory-attended* condition. We used a 2*2 (RC × Group) circular ANOVA (Bonferroni corrected for multiple comparisons) to examine whether the 3-Hz phase in the *Auditory-attended* condition differed as a function of attention abilities.

## Supplementary Figures

**Figure S1: Component-Specific Differences in Entrainment Strength**

To assess attention modulation of component-space neural entrainment strength, for each experiment, we conducted a 2×2 repeated-measures ANOVA (RC × Condition) on the 3-Hz amplitude extracted from component-space data.

In **Experiment 1**, results showed a significant main effect of RC such that the overall 3-Hz amplitude was higher for RC1 compared to RC2 [F_(1, 42)_ = 98.42, p <.001]. No other significant effects were observed for condition [F_(1, 42)_ = 0.174, p =.679] or interaction [F_(1, 42)_ = 1.68, p =.2]. Similar to Experiment 1, in **Experiment 2**, results reveal a significant main effect of RC with a higher 3-Hz amplitude for RC1 compared to RC2 [F_(1, 71)_ = 73.01, p <.001]. No other significant effects were observed for condition [F_(1, 71)_ = 1.38, p =.244] or interaction [F_(1,71)_ = 0.002, p =.98]. See Figure S1 for a full results description.

**Figure S1:**
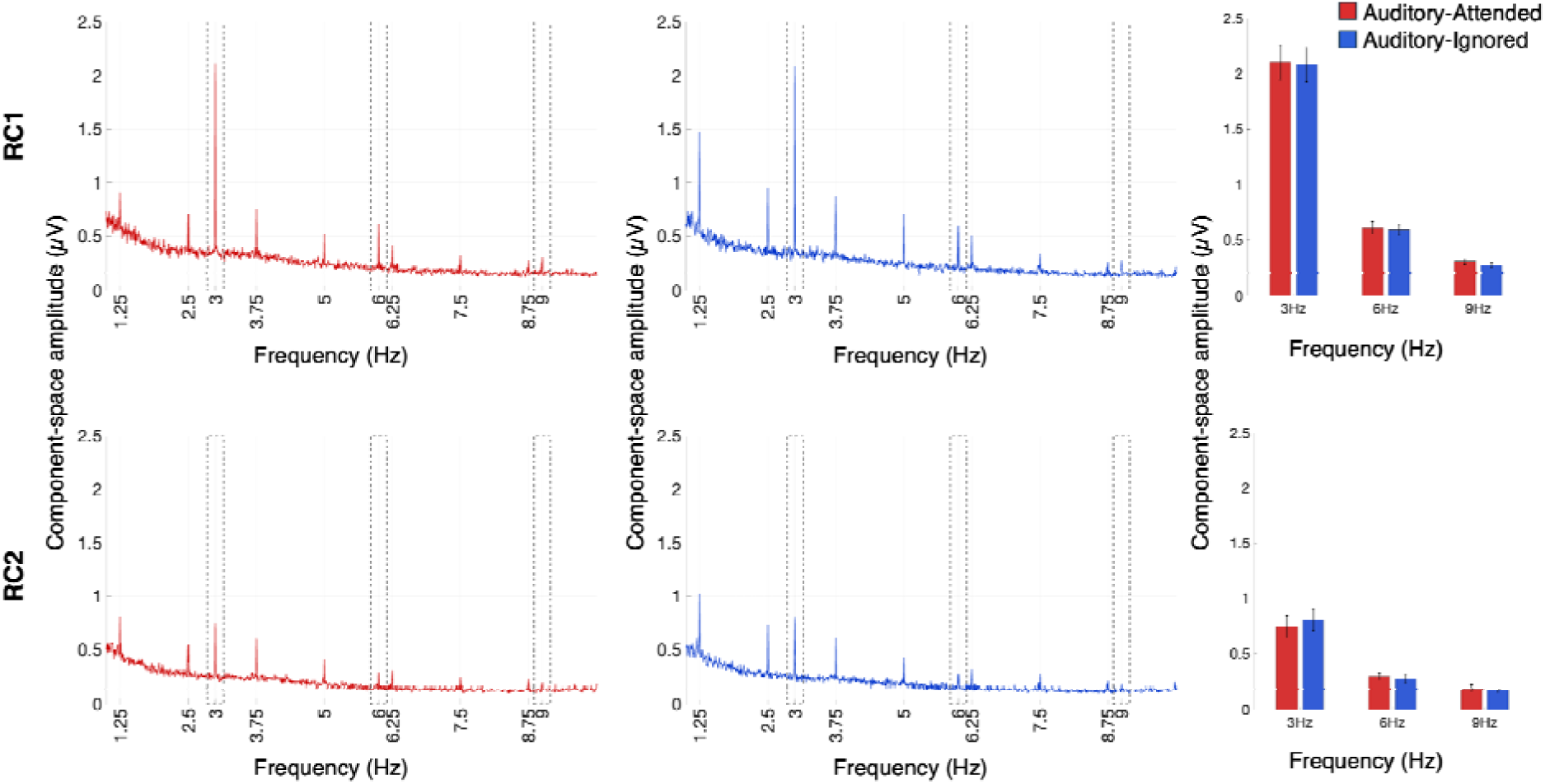
Power spectrum of the component-space data showing the 3-Hz (and harmonics) amplitude across experiments, conditions, and components. Top panels: data from Experiment 1; Bottom panels: data from Experiment 2. (A) Full power spectrum for Auditory-attended (red) and Auditory-ignored (blue). Peaks are aligned with frequencies of the auditory and the visual stimuli (3-Hz and 1.25-Hz, and harmonics, respectively). Dashed rectangles highlight the frequencies of the auditory stimuli and harmonics. (B) Bar graphs of the mean values of 3-Hz frequency amplitude and its harmonics. Vertical lines represent standard errors. The top panels represent RC1 component space data, and the bottom panels represent RC2.

**Table S1:** Demographic and Clinical Characteristics of the Sample.

| Variable | Category | Experiment 1 (n=43) |  | Experiment 2 (n=36) |  |
| --- | --- | --- | --- | --- | --- |
|  |  | n | % | n | % |
| Gender | Male | 21 | 48.8 | 23 | 63.9 |
|  | Female | 22 | 51.2 | 13 | 36.1 |
| Ethnicity | White | 26 | 60.4 | 22 | 61.1 |
|  | Asian | 18 | 41.8 | 11 | 30.5 |
|  | Other | 1 | 2.3 | 7 | 9.4 |
| Handedness | Right | 37 | 86 | 33 | 94.4 |
|  | Left | 6 | 14 | 3 | 5.6 |
| Primary Language | English | 30 | 69.7 | 26 | 72.2 |
|  | Other | 13 | 30.3 | 10 | 27.8 |
| Former Diagnosis | ADHD | 5 | 11.6 | 10 | 27.7 |
|  | Other learning difficulties | 7 | 16.2 | 14 | 38.8 |
| Medication Type | Stimulant | 2 | 4.6 | 5 | 13.8% |
|  | Antihistamine | 2 | 4.6 | 1 | 2.8% |
|  | Anti-inflammatory | 1 | 2.3 | 2 | 5.6% |
|  | Mood stabilizers | 0 | 0% | 1 | 2.8% |
|  | Other | 1 | 2.3 | 1 | 2.8% |
|  | No medication use | 37 | 86 | 37 | 86% |

*Values represent counts (n) and percentages (%). ADHD = attention-deficit/hyperactivity disorder.*

**Experiment 1:** A total of 43 participants were included in the final analysis of the first experiment (age range: 10–12 years, M = 10.8, SD = 0.72). The sample comprised 21 males (48.8%) and 22 females (51.2%). Most participants were right-handed (86.0%), and English was the primary language for 30 participants (69.7%), while 13 participants (30.3%) reported being bilingual or speaking a different primary language. In terms of ethnicity, 26 (60.4%) identified as White, 18 (41.8%) as Asian, and 1 (2.3%) as American Indian. A prior diagnosis of attention-deficit/hyperactivity disorder (ADHD) was reported by 5 (11.6%) participants, while 7 (16.2%) indicated a specific learning difficulty, such as dyslexia. Regarding medication, 6 (13.9%) participants were currently taking prescribed medication as detailed in the table.

**Experiment 2:** A total of 36 participants were included in the final analysis of the first experiment (age range: 12–14 years, M = 12.9, SD = 0.6). The sample comprised 23 males (63.9%) and 13 females (36.1%). Most participants were right-handed (94.4%), and English was the primary language for 26 participants (72.2%), while 10 participants (27.7%) reported being bilingual or speaking a different primary language. In terms of ethnicity, 22 (61.1%) identified as White, 11 (30.5%) as Asian, and 7 (9.4%) as multiracial. A prior diagnosis of attention-deficit/hyperactivity disorder (ADHD) was reported by 10 (27.7%) participants, while 14 (38.8%) indicated a specific learning difficulty, such as dyslexia. Regarding medication, 9 (25%) participants were currently taking prescribed medication.

## Footnotes

1 https://github.com/dmochow/rca

