## Supplementary figures and images for "Auditory attention reorganizes the phase alignment of neural oscillations"

### Supplemental Figure 1

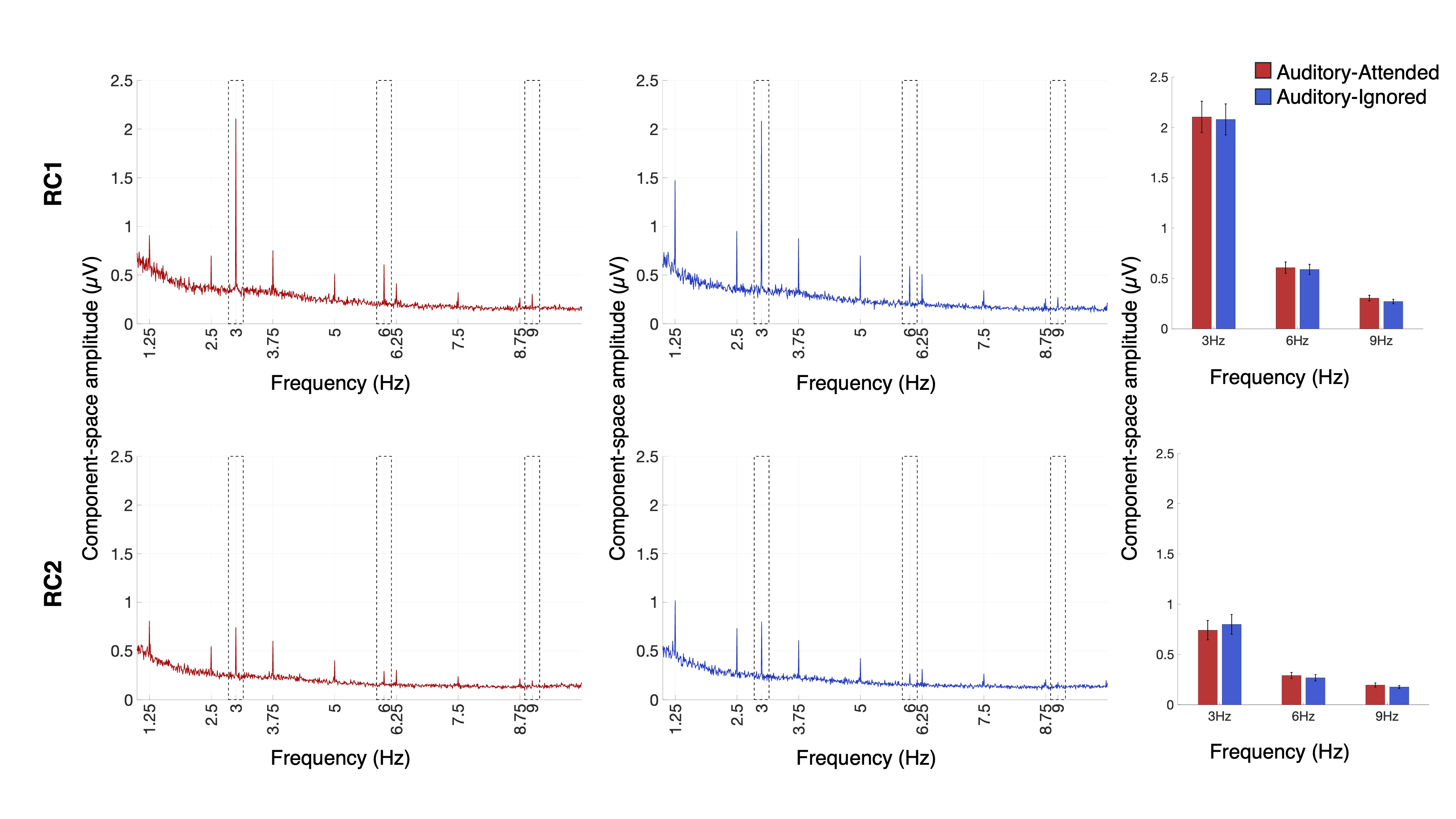
